# Inter-subject phase synchronization and the dynamics of human cognition

**DOI:** 10.1101/167072

**Authors:** Taylor Bolt, Jason S. Nomi, Shruti G. Vij, Catie Chang, Lucina Q. Uddin

## Abstract

Massive whole-brain blood-oxygen-level dependent (BOLD) signal modulation (up to 95% of brain voxels) in response to task stimuli has recently been reported in functional MRI investigations. These findings have two implications. First, they highlight inability of a conventional ‘top-down’ general linear model approach to capture all forms of task-driven brain activity. Second, as opposed to a static ‘active’ or ‘non-active’ localization theory of the neural implementation of cognitive processes, functional neuroimaging should develop and pursue *dynamical* theories of cognition involving the dynamic interactions of *all* brain networks, in line with psychological constructionist theories of cognition. In this study, we describe a novel exploratory, bottom-up approach that directly estimates task-driven brain activity regardless of whether it follows an *a priori* reference function. Leveraging the property that task-driven brain activity is associated with reductions in BOLD signal variability, we combine the tools of instantaneous phase synchronization and independent component analysis to characterize *whole-brain* task-driven activity in terms of group-wise similarity in temporal signal dynamics of brain networks. We applied this novel framework to task fMRI data from a motor, theory of mind and working memory task provided through the Human Connectome Project. We discovered a large number of brain networks that dynamically synchronized to various features of the task scan, some overlapping with areas identified as ‘active’ in the top-down GLM approach. Using the results provided through this novel approach, we provide a more comprehensive description of cognitive processes whereby task-related brain activity is not restricted to dichotomous ‘active’ or ‘non-active’ inferences, but is characterized by the temporal dynamics of brain networks across time.

**Significance Statement:** This study describes the results of a novel exploratory methodological approach that allows for direct estimation of task-driven brain activity in terms of group-wise similarity in temporal signal dynamics, as opposed to the conventional approach of identifying task-driven brain activity with a hypothesized temporal pattern. This approach applied to three different task paradigms yielded novel insights into the brain activity associated with these tasks in terms of time-varying, low-frequency dynamics of replicable synchronization networks. We suggest that this exploratory methodological approach provides a framework in which the complexity and dynamics of the neural mechanisms underlying cognitive processes can be captured more comprehensively.

## Introduction

Task-based fMRI has produced a wealth of large-scale function-structure mappings representing associations between mental processes and unique regions or networks of the brain^1–5^. However, a consensus localization of mental functions to unique cortical/sub-cortical areas has yet to emerge. Recent findings reminiscent of early theories of cortical mass action^6,7^ have demonstrated that time-locked activation is observed in a majority of the brain (in some cases, over 95% of the brain) in response to external task presentations^8–12^. For example, using massive within-subject averaging, Gonzalez-Castillo et al.^8^ observed that the majority of the brain exhibits time-locked activation to a simple visual stimulation and attention task, with substantial variation of hemodynamic responses across different regions of the brain.

These results suggest that it is not a question of ‘if’ most brain regions are involved in any given task, but rather ‘when.’ These results further motivate a dynamical theory of cognition that goes beyond dichotomous decisions of ‘active’ or ‘non-active’ brain areas. Rather, an adequate description of psychological processes involves identifying the temporal dynamics of brain networks periodically synchronized to external task-relevant events of the task presentation. A constructionist account of cognitive processes^13,14^ provides a theoretical grounding for such a theory whereby cognition emerges from the interaction and temporal dynamics of more basic domain-general, functional brain networks (e.g. fronto-parietal network, default-mode network, etc.). Therefore, an adequate description of the cognitive processes involved in an experimental task requires examination of all brain networks that periodically synchronize (simultaneously or consecutively) to various task features.

In typical fMRI cognitive neuroscience paradigms, a brain region is understood to be involved in a task if that region exhibits blood-oxygen-level dependent (BOLD) variation that follows a hypothesized temporal structure. This ‘top-down’ approach often involves the application of a general linear model (GLM), and a reference function corresponding to task events that is convolved with a hemodynamic response function (HRF)^15–17^. However, any brain regions or networks involved in a given cognitive process not predicted by conventional reference functions would not be detected by this approach. Most importantly, the ‘top-down’ approach would not capture complex temporal dynamics that a given task may produce in activation patterns across the brain.

We propose a ‘bottom-up’ exploratory approach for the analysis of task fMRI data that directly estimates task-driven brain activity in a data-driven manner, without the application of a hypothesized temporal structure. The estimation of task-driven brain activity is non-trivial, as brain activity is highly dynamic and structured even in the absence of stimulus presentations, owing to spontaneous activity^18–21^ and signal fluctuations due to noise sources^22,23^. However, a potential marker of task-driven brain activity (opposed to spontaneous activity or noise) is *variability reduction* in the neural signal post-stimulus. A reliable property of task-driven BOLD activity^24,25^, and firing rates and membrane potentials from single-cell recordings^26,27^, is post-stimulus signal variability reduction. An approach incorporating the neural variability reduction phenomenon could be used as an exploratory measure of task-driven brain activity. Established inter-subject synchronization measures provide the foundation necessary for detecting task-related signal variability reductions^28–32^, assuming a brain region is entrained to a stimulus in a common way *across participants*. The logic of this approach is as follows: in individuals concurrently experiencing the same stimulus, a task-responsive brain region entrained to that stimulus would produce similar or conserved temporal dynamics across subjects^29^ as a result of the common entrainment of that voxel to the stimulus condition.

Conventional ‘static’ inter-subject correlation approaches^30,31^ provide an average measure of brain synchrony across the entire task at each voxel. For a dynamical measure of inter-subject synchronization, the current paper utilizes an instantaneous phase synchronization measure^28,32^ for voxel-wise assessments of synchronization at each time point (**Figure 1**), followed by an independent components analysis (ICA)^33,34^ applied to the voxel-wise synchronization time series. This novel methodology identifies time-varying, task-driven brain network dynamics without dependence on *a priori* reference functions, and provides a framework for the examination of task-relevant *whole-brain temporal dynamics*.

**Figure 1.**
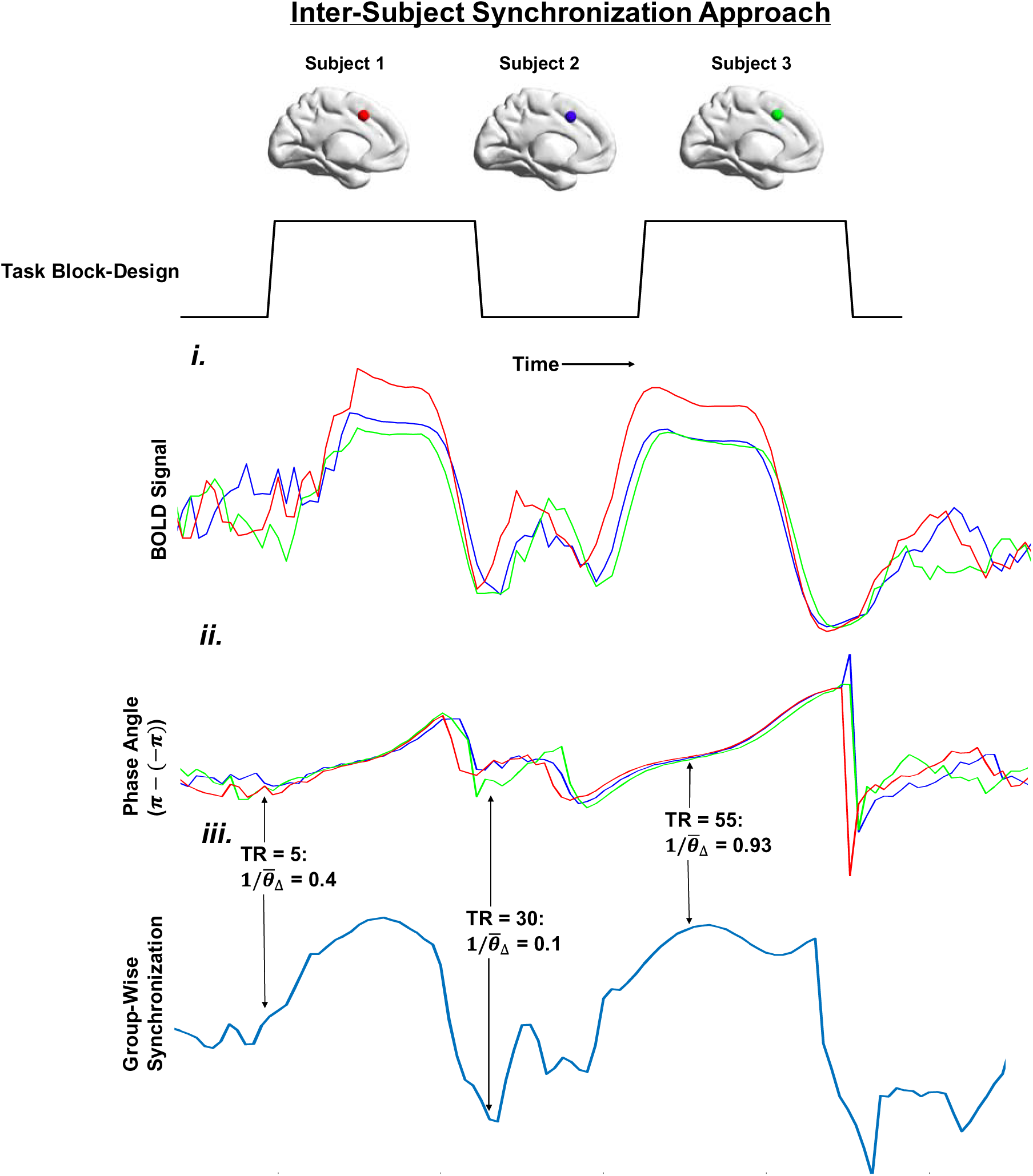
Using inter-subject synchronization to study task-driven brain responses. If one assumes that a task-responsive brain region (e.g. dorsal anterior cingulate cortex; dACC) exhibits a common response *across participants* (e.g. increase in signal amplitude), one can implement an inter-subject synchronization approach that can examine that brain region’s across-subject variability at each time point by measuring synchronization of the BOLD signal across participants at that time point. **I**) Three subject’s dACC time series are displayed in separate colors, and all demonstrate a similar increase in signal amplitude in the task-blocks. **II**) In order to focus our analysis on ‘pure’ *temporal* structure across subjects (as opposed to magnitude differences across subjects), a phase representation of each subject’s time series is constructed^28^. **III**) From the phase representation of each signal, we can identify group-wise synchronization estimates for each time point by essentially calculating the (inverse) average angular distance between each subject’s phase signal. Using group-wise synchronization estimates at each time point, we can examine synchronization dynamics across the task scan in a data-driven manner, without any reference to a hypothesized on- and off-task block structure or HRF specification. Phase calculations demonstrate higher phase synchrony during a task block (time point = 55) compared to non-task blocks (time point = 5 and time point = 30).

There are three methodological and theoretical advantages of this methodology over the traditional GLM approach: 1) a reference function of task events or hemodynamic response function (HRF) is not required to detect task-responsive brain regions, 2) the synchronization approach does not assume that task-driven brain responses follow a single, simple form (e.g. transient, sustained or mixed activation) of voxel-wise activity, and 3) the approach allows for assessing dynamical or time-varying task-related brain networks. To demonstrate the viability and benefits of this novel methodology when studying task-driven brain responses, we applied this approach to three large-sample task Human Connectome Project (HCP)^35^ fMRI data sets: motor, theory of mind, and working-memory.

The results demonstrate that this alternative methodological approach recapitulates traditional GLM activation maps, but more importantly, it also identifies novel time-varying replicable synchronization networks. The identification of these new novel synchronization networks, in addition to brain areas commonly identified by a traditional GLM approach, provides a richer, fuller description of cognition whereby task-related brain activity is not restricted to dichotomous ‘active’ or ‘non-active’ inferences, but rather is more comprehensively described by the temporal dynamics of brain networks across time.

## Results

### Instantaneous Phase Analysis

As phase analysis is most effective on a band-limited signal^28^, a data-driven estimation of the frequency band for all three tasks was conducted to guide the selection of a frequency band of interest. A static inter-subject synchrony analysis (see *Experimental Procedures*) revealed that a low frequency sub-band (~0.01 - 0.085 Hz) demonstrated the highest level of average BOLD synchrony across subjects. This sub-band was used for further analyses (**Figure S1**). The average BOLD synchrony maps of this sub-band for each task overlapped significantly with the z-scored ‘activation’ maps derived from the GLM reference-function approach (*r_Motor_* = 0.76, *r_TOM_* = 0.85; *r_WM_* = 0.83; **S1 Figure**).

Next, instantaneous phase synchronization analysis^28^ was conducted on this low-frequency sub-band, providing group-wise synchronization time-course estimates at each voxel. Then, an ICA was applied to the voxel-by-voxel synchronization time series to parse synchronization dynamics into distinct spatial synchronization networks. A range model-order ICA solutions were estimated (10, 15, 20 components) as there was no *a priori* indication of the optimal number of synchronization networks relevant to each task. To guide the ICA model-order solution for each task, an ICA was performed on a separate confirmatory sample of 75 subjects in which the same synchronization analysis was applied. Additionally, this analysis provided an assessment of the robustness of the synchronization approach to sampling characteristics. The optimal solution for each task maximized the replicability between the original and confirmatory samples, as measured by the number of matching components, the spatial correlation between matching ‘non-noise’ components, and the temporal correlation between matching ‘non-noise’ component time series. For all tasks, all ICA solutions yielded satisfactory replication across samples, but the 10-component solution yielded the highest replicability according to all three metrics (Motor/TOM/WM: *matching components*: 10/9/10 out of 10; *r̄*_Spatial_: 0.80/0.68/0.80, *r̄*_Temporal_: 0.96/0.89/0.93) compared to the 15 component (*matching components*: 15/12/15 out of 15; *r̄*_Spatial_: 0.7/0.58/0.69, *r̄*_Temporal_: 0.90/0.88/0.87) and 20 component solutions (*matching components*: 17/16/18 out of 20; *r̄*_Spatial_: 0.68/0.57/0.65, *r̄*_Temporal_: 0.92/0.87/0.91). Results from the 10-component solution are thus presented here.

#### Synchronization Dynamics During the Motor Task

The motor task was chosen because the neural correlates of movement are well understood^36^, providing a validation test for the synchronization approach. The activation maps for each motor movement produced by the standard GLM-reference function approach corresponded precisely to their expected task-relevant brain areas (**Figure 2**). For example, left motor cortex activation was associated with right-hand motor movements while fronto-parietal and visual activation was associated with visual cue trials.

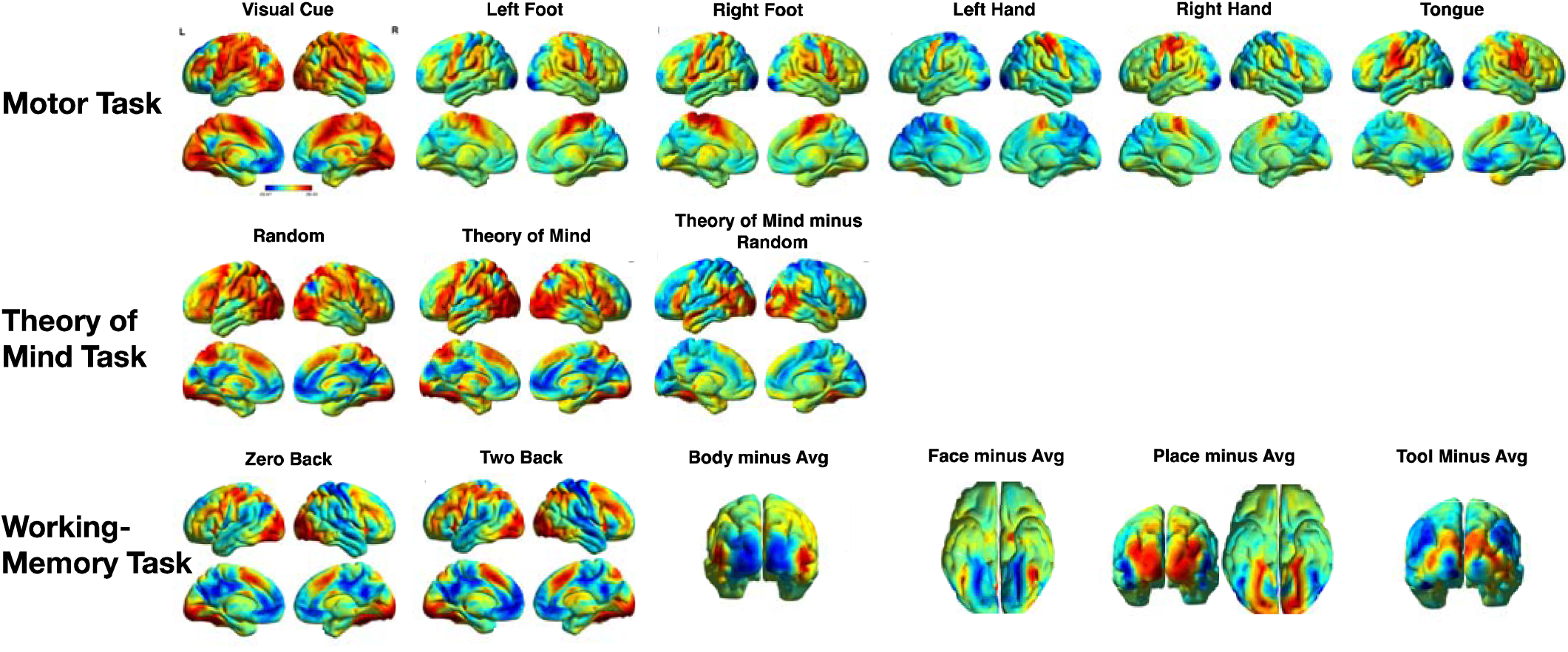

between two task-conditions (i.e. subtraction contrasts).

Three of the ten components derived from the ICA synchronization analysis were classified as noise (**S3 Figure**) as they had many voxels with strong weights in either white matter, cerebrospinal fluid (CSF) or along the surface of the brain corresponding to movement artifacts. All seven non-noise components showed a characteristic pattern of peaks and troughs (i.e., local maxima and minima) of synchronization across the course of the scan with varying degrees of periodicity (**Figure 3**). Significant synchronization peaks were defined as those collection of synchronization time points with estimates greater than a significance threshold (represented as the horizontal dotted lines superimposed on synchronization time-courses in each figure), determined through the application of a bootstrap resampling procedure. Interestingly, the first five component synchronization time series were uniquely time locked to the five different movement blocks (C1 – Tongue: *r* = 0.7; C2 – Left Hand: *r* = 0.62; C3 – Right Foot: *r* = 0.66; C4 – Right Hand: *r* = 0.6; C5 – Left Foot: *r* = 0.59; *p’s* < 0.0001). Additionally, the spatial patterns of the first five components corresponded precisely to their expected representation in the motor cortex (contralateral) and cerebellum (ipsilateral). For example, C1 (tongue) consisted of the bilateral ventrolateral motor cortex while C2 (left hand) consisted of the right dorsolateral motor cortex and left ipsilateral cerebellum (**Figure 3**) corresponding to the tongue and hand areas of the motor cortex respectively. These results are consistent with the results from the standard GLM approach, but more importantly, motor function localization was achieved without *a priori* specification of a reference function or other features of the standard GLM approach.

**Figure 3.**
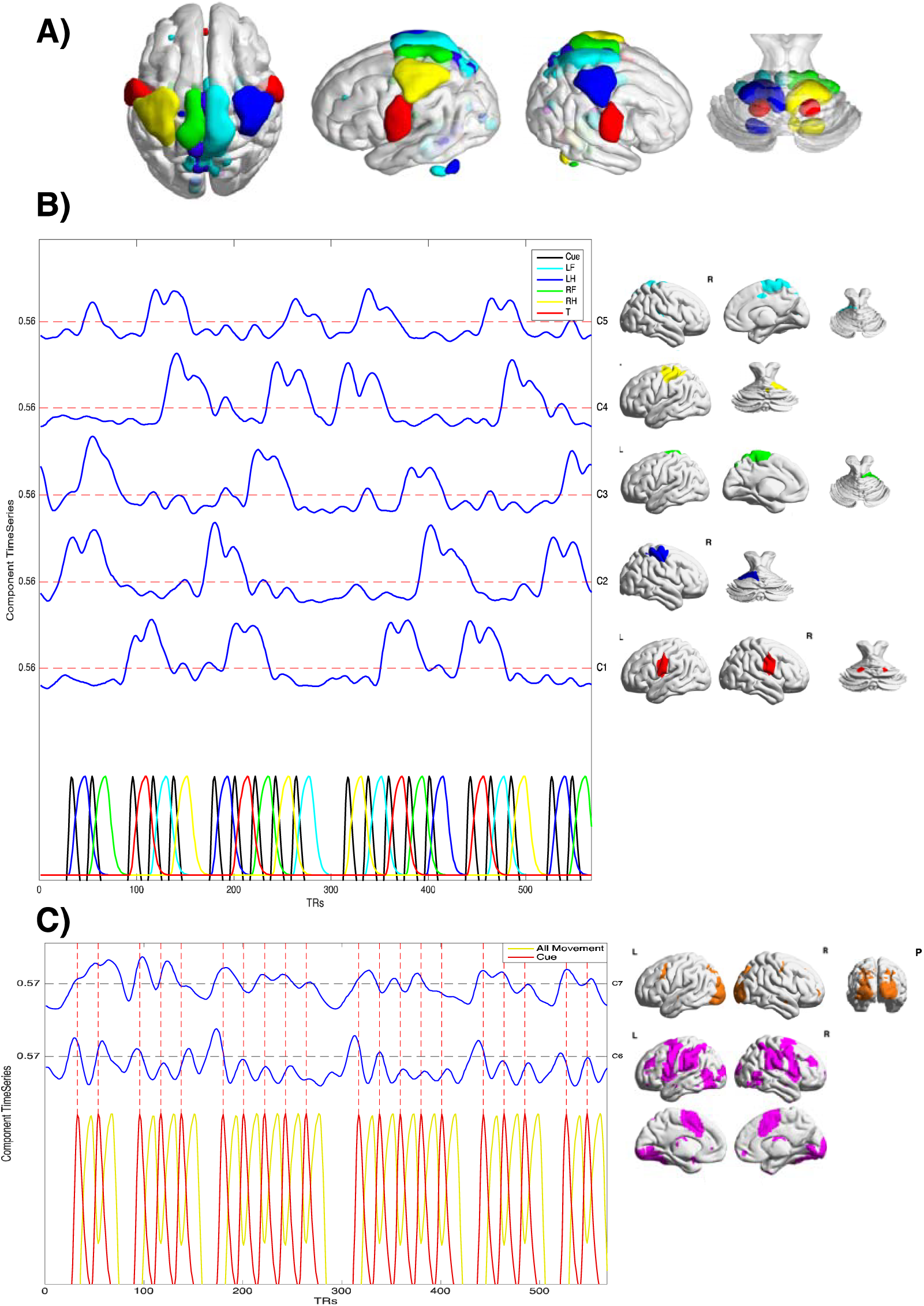
Synchronization Dynamics of Motor Components. (C = Component; R = Right; L = Left; Cue = Visual Cue; LF = Left Foot; LH = Left Hand; RF = Right Foot; RH = Right Hand; T = Tongue). **A)** Cortical and cerebellar ROI-volume drawings of the thresholded spatial weights for C1 through C5 (colors are the same as the surface visualizations below; visualized with BrainNet Viewer^37^. ICA on the group-wise synchronization estimates achieved a precise localization of the five components, corresponding to the five different movement types, along their corresponding representation in the motor and cerebellar cortices. **B)** The synchronization time courses for C1 through C5 with its corresponding spatial surface representation (same color as volume drawing above), and the GLM reference functions with colors corresponding to the color of the component it is most associated with. Significance thresholds for each synchronization time course are represented by the red dotted-lines (with the corresponding value to the left of the plot). Synchronization values for each time course varies from 0 to 1. **C)** The synchronization time courses and corresponding thresholded spatial weights of C6 (magenta) and C7 (orange), and the GLM reference functions (visual cue in red, and all movement types together in yellow). For visual reference, the peaks of the reference function corresponding to the visual cue trials are plotted vertically with a red dotted-line. Significance thresholds for each synchronization time course are represented by the horizontal black dotted-lines (with the corresponding value to the left of the plot).

The results for the visual cue with the standard GLM-reference function approach revealed strong activation in voxels that encompassed both visual and fronto-parietal control areas (Figure 2). However, the novel synchronization approach revealed differing temporal dynamics between visual and fronto-parietal control areas elucidating the temporal interplay between these two brain areas. Significant peaks in the synchronization time course of C6 (fronto-parietal control areas) and C7 (visual areas) occur predominantly around visual-cue onset indicating the start of motor movement trials (Figure 3). C7 was most strongly correlated with visual cue onset (*r* = 0.45, *p* < 0.0001), consistent with the strong spatial weights in the visual cortices for this component. The synchronization time series of C6 was moderately positively correlated with the visual cue blocks (*r* = 0.26, *p* < 0.0001) and strongly negatively correlated with all motor blocks (*r* = −0.58, *p* < 0.0001; correlated with a single combined convolved regressor with all motor block types). Significant synchronization peaks for C6 occurred predominantly at the beginning of one of the six movement block sequences (occurring after each of the six 15-second fixation blocks), perhaps corresponding to the allocation of attentional resources to the onset of a new sequence of task demands after a block of fixation. Interestingly, the spatial pattern of C6 also includes the entire motor cortex, possibly representing a ‘preparatory’ motor signal for the onset of the motor blocks.

#### Synchronization Dynamics During the Theory of Mind Task

The theory of mind task consisted of short video clips of either interacting or randomly moving shapes, and is used to study higher-level processing of dynamic visual stimuli. The activation maps produced by the standard GLM reference-function approach were strongly similar between the theory of mind and random blocks (**Figure 2**). Activation for both block types was observed in primary and higher-order visual cortices, and fronto-parietal areas. Activation differences between the theory of mind and random blocks were primarily observed in lateral occipital cortices, lateral temporal-parietal junction, and inferior frontal gyri.

Two of the 10 components derived from the ICA phase synchronization analysis were classified as noise while one component did not replicate in the confirmatory sample and was not interpreted further (**S3 Figure).** Four of the seven non-noise components were observed to have strong spatial weights predominantly in the primary and higher-order visual cortices (C2, C4, C5, and C7; **Figure 4**), perhaps related to the fact that each task block required careful visual attention to moving task stimuli. The other components (C1, C3 and C6) were observed to have strong spatial weights predominantly in areas of the fronto-parietal control network (C1 and C6) and temporal lobes (C3). The synchronization dynamics of the seven signal components can be roughly divided into off-task-block (C2, C5 and C7), on-task-block (C1 and C6), task-block specific (C3) and non-block (C4) components.

**Figure 4.**
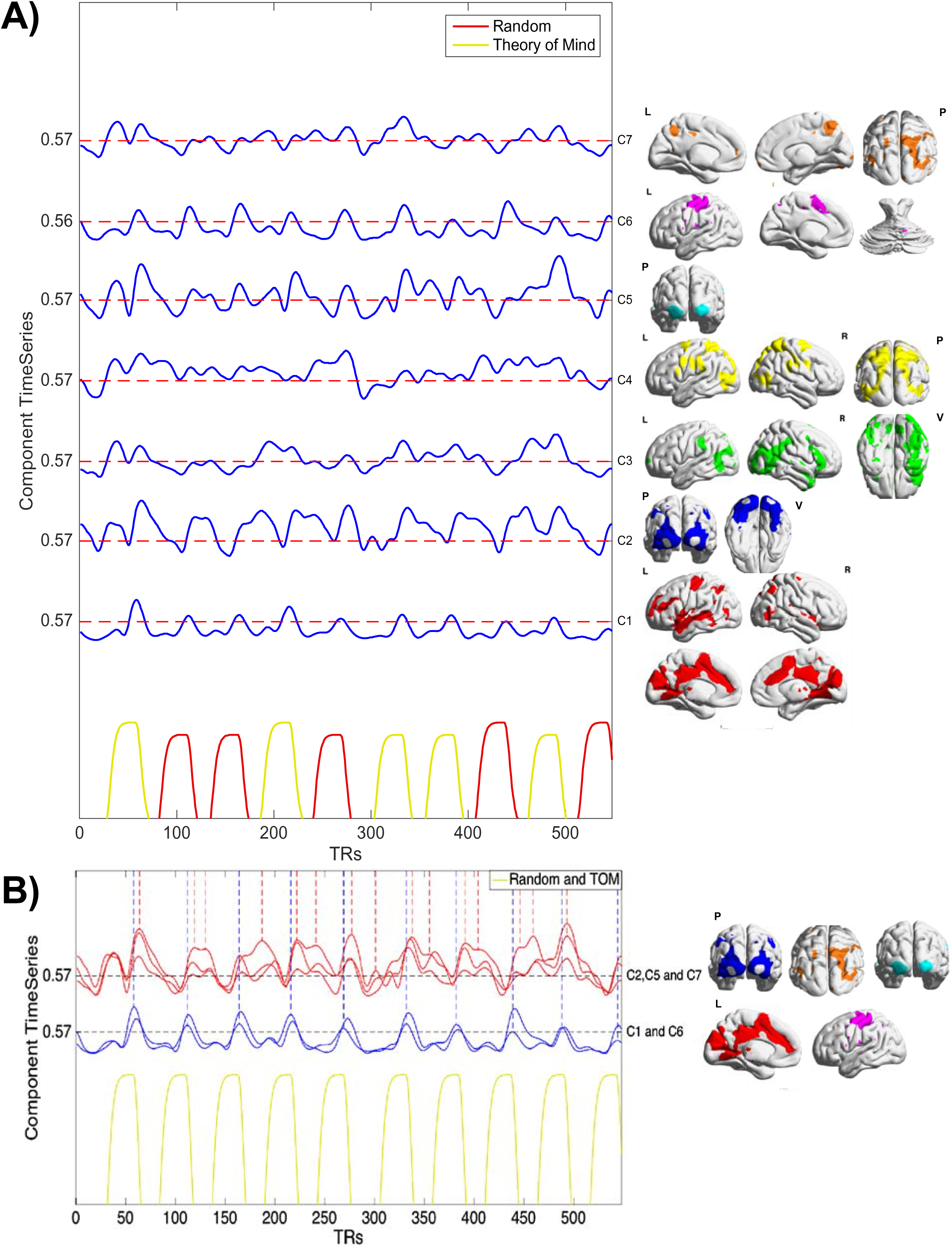
Synchronization Dynamics of Theory of Mind Components. (L = Left; R = Right; P = Posterior; V = Ventral; TOM = Theory of Mind). **A)** The synchronization time courses and corresponding thresholded spatial weights of C1 through C7, and the GLM reference functions (random shape movement in red, and intentional shape movement in yellow) for the task. Thresholds for each synchronization time course are represented by the red dotted-lines (with the corresponding value to the left of the plot). **B)** Visual comparison of the on- and off-task block component synchronization time courses (both block types are combined in one reference function and are presented in yellow). On-block task component (C1 and C6) synchronization time courses are presented in blue and observed to have significant synchronization peaks during the last seconds of the 20s task-blocks extending into the off-block periods. Off-block task component (C2, C5 and C7) synchronization time courses are presented in red, and are observed to have significant synchronization peaks shortly after the on-black task component peaks during the off-block periods. For visual reference, statistically significant peaks in the synchronization time courses for C1 (on-block component) and C2 (off-block component) are marked by vertical blue- and red-dotted lines. Significance thresholds for the synchronization time courses are represented by the black dotted-lines (the threshold for C6 was 0.56, but presented as 0.57 for visualization purposes).

C3 was classified as a task-block specific component because its synchronization time series positively correlated with theory of mind block types (*r* = 0.71, *p* < 0.0001). Additionally, the C3 spatial weights within the bilateral fusiform gyri and right tempo-parietal junction (TPJ) correspond to brain areas consistently implicated in the theory of mind literature^38,39^. Interestingly, the spatial weights of C3 significantly overlapped (*r* = 0.41, *p* < 0.0001) with the theory of mind-minus-random contrast (**Figure 2**), representing the relative activation between the theory of mind and random condition. Spatial weights for the non-block component, C4, corresponded to posterior parietal areas and area V5 (visual motion perception) while the synchronization time course of C4 was observed to have statistically significant synchronization peaks across both task runs of the theory of mind task. This indicated that this network may be relevant for domain-general visual attention of moving stimuli.

C1 and C6, corresponding to dACC/posterior cingulate cortex (PCC)/left temporal lobe and supplementary motor cortex/left motor cortex, respectively, were classified as on-task block components because the synchronization time courses were positively correlated with all task blocks (C1: *r* = 0.5, *p* < 0.0001; C6: *r* = 0.2, *p* < 0.0001), and synchronization peaks of both component time courses occurred during task-blocks. C2, C5 and C7, corresponding to bi-lateral secondary visual cortices, right lateral occipital/posterior parietal cortices, and primary visual cortices, respectively, were classified as off-block components because the synchronization time series of the components were not positively correlated with task blocks (C2: *r* = −0.4, *p* < 0.0001; C5: *r* = - 0.04, *p* = 0.396; C7: *r* = −0.03, *p* = 0.485), and synchronization peaks of the component time courses occurred during off-block components. Interestingly, the five on- and off-task block components showed a characteristic pattern of synchronization dynamics across the duration of the theory of mind task (**Figure 4**). Specifically, the on-task block components were observed to have significant synchronization peaks during the last seconds of the 20s task-blocks extending into the off-block periods, with subsequent synchronization peaks of the off-task block components, in the off-block periods. The onset of cognitive processing associated with C1, and subsequent motor processing associated with C6 (mean TR difference between synchronization peaks of C1 and C6: 2.01 TRs, *SD*: 0.88) towards the latter end of the task-block is consistent with the fact that after the 20s on-block video watching period, the participants were asked to rate the video in terms of intentional or randomly acting shapes. During the rating period and subsequent fixation period, participants were presented with static visual stimuli, consistent with the onset of synchronization peaks of the different visual components C2, C5, and C7.

#### Synchronization Dynamics During the Working-Memory Task

The working-memory task was a visual N-back task with 0-back and 2-back conditions consisting of category-specific visual stimuli. The activation maps for the 0-back and 2-back conditions produced by the standard GLM-reference function approach were similar, with the strongest activation values in areas of the fronto-parietal control network (**Figure 2**). Differences in activation associated with each category-specific stimulus (body, face, place, and tool stimuli) were observed in the primary visual and inferior-temporal cortices.

One of the ten components derived from the ICA phase synchronization analysis was classified as noise (**S3 Figure**). Four non-noise components (**Figure 5**) were grouped together because they had strong spatial weights in visual cortices and similar synchronization time courses. The spatial weights of C1 through C4 demonstrated that each component represented a unique partition of primary and secondary visual cortices. The spatial weights for the four components overlapped to a moderate degree with the differential activation areas associated with each category-specific stimuli: C2 was spatially correlated with body stimuli activation (*r* = 0.49, *p* < 0.0001), C1 and C4 were spatially correlated with place stimuli activation (C1: *r* = 0.51; C4: *r* = 0.43; *p*’s < 0.0001), C3 was spatially correlated with tool stimuli activation (*r* = 0.43; *p* < 0.0001), and C4 was not strongly correlated with any of the four stimuli.

**Figure 5.**
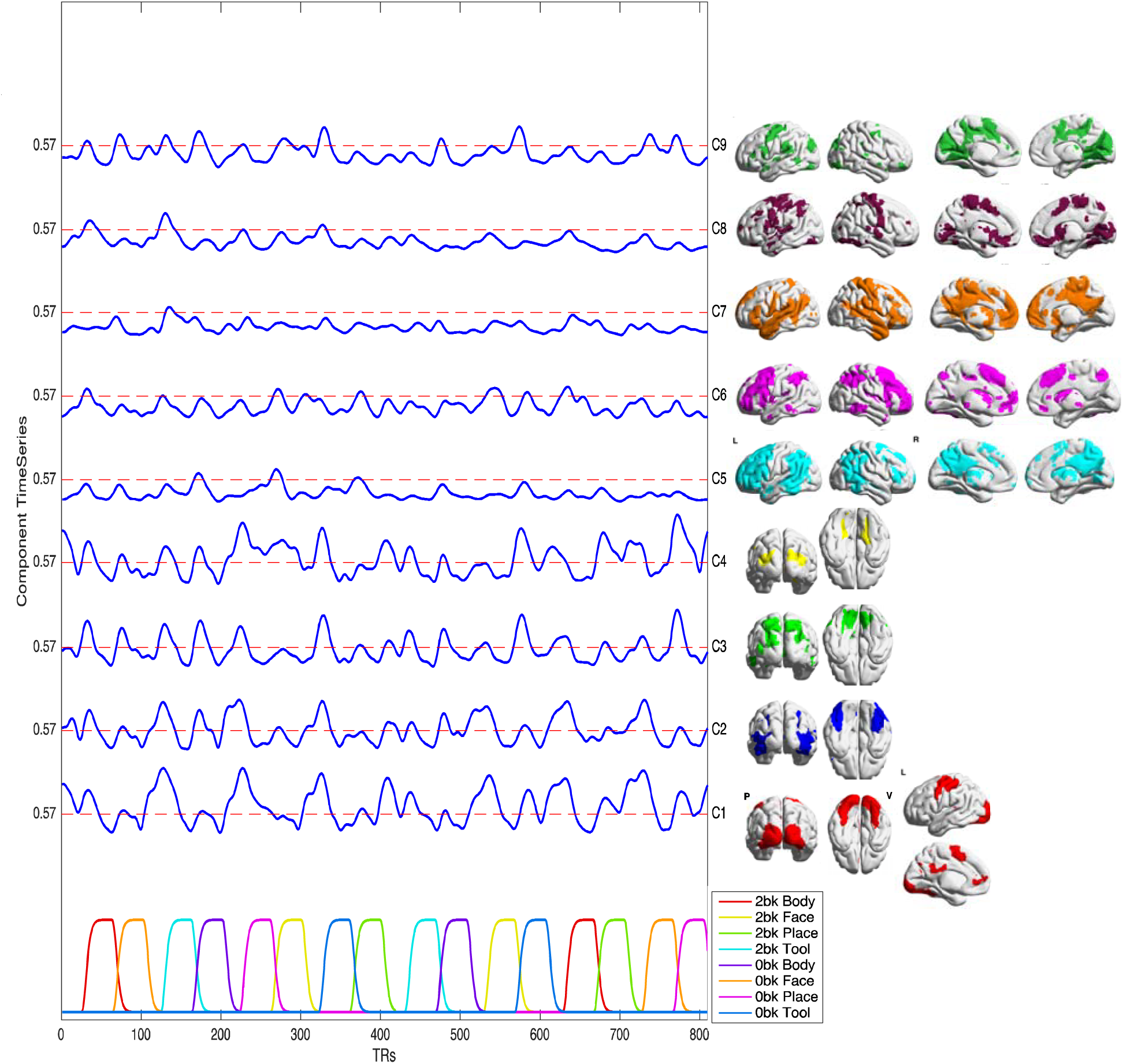
Synchronization Dynamics of Working-Memory Components. (L = Left; R = Right; P = Posterior; V = Ventral). The synchronization time courses and corresponding thresholded spatial weights of C1 through C9, and the GLM reference functions (2-Back and 0-Back conditions with four category-specific stimuli,) for the task. Thresholds for each synchronization time course are represented by the red dotted-lines (with the corresponding value to the left of the plot).

While the subtraction contrasts between the stimulus blocks revealed differential activation patterns in visual cortices, the phase synchronization approach revealed that visual areas, represented by components C1-C4, demonstrate predominantly similar temporal dynamics to all types of stimulus blocks, with significant synchronization peaks occurring at approximately the same time points, but with differences in the degree of synchronization at each peak time point. In relation to the reference functions provided in the task, the synchronization time courses for all four components contained approximately 16 significant synchronization peaks, corresponding to the 16 task blocks of all types presented to the participants in the working-memory task. However, increases in synchronization estimates did not coincide with task blocks, but were negatively correlated with task blocks (C1: *r* = −0.68; C2: *r* = −0.65; C3: *r* = −0.32; C4: *r* = −0.26; *p’s* < 0.0001), with significant synchronization peaks occurring primarily in the off-block periods, sometimes beginning around the latter end of task blocks. Four unique combinations of the four components were preferentially negatively associated with the four category-specific stimuli types (in one case, for C4, positively correlated): C1 and C4 with body stimuli (C1: *r* = −0.36; C4: *r* = −0.34; *p’s* < 0.0001), C1 and C2 with face stimuli (C1: *r* = −0.24; C2: *r* = −0.25; *p’s* < 0.0001), C2, C3 and C4 with place stimuli (C2: *r* = −0.26; C3: *r* = −0.17; C4: *r* = 0.31; *p’s* < 0.0001), and C1, C3, and C4 for tool stimuli (C1: *r* = −0.15; C2: *r* = −0.18; C4: *r* = −0.18, *p’s* < 0.0001). The negative correlation between the four component synchronization time courses with task blocks suggests that these components are related to the eight 15-second visual fixation blocks between some task-block periods. However, the fact that approximately 16 significant synchronization peaks were observed across the task scan, corresponding to the number of task blocks, suggests that peaks of the four components correspond to synchronized downturns in the BOLD response to the task blocks that occur primarily toward the end of each task block. This interpretation was confirmed in comparison of the component average BOLD time courses with the synchronization time courses (**S5 Figure/Discussion**).

Other interesting findings pertain to the other five non-noise components (**Figure 5**). Examination of the synchronization time courses of C5, C7, and C8 reveals that these components exhibit significant synchronization peaks early in the working-memory task, but diminished, non-significant synchronization peaks thereafter, indicating habituation of these networks to the working-memory task after repeated task-block presentations, or their involvement in task-set initiation. The spatial patterns of C5, C7 and C8 suggest these components correspond to the posterior parietal and left-dominant prefrontal brain areas, default mode network, and bi-lateral motor cortex/basal ganglia areas, respectively. Of note, in addition to strong spatial weights for C5 in lateral posterior parietal and prefrontal areas, they were also observed in the precuneus, a default-mode network hub^40^, indicating that default-mode network regions may synchronize with executive control regions during certain time points of the working-memory task^41,42^. Strong spatial weights for C7 were also observed in the anterior caudate nucleus, an area that forms a central part of a cortico-striatal loop with frontal regions and known to be important in cognitive processing^43,44^. Spatial weights of C6 and C9 corresponded to task-positive, executive-control brain areas and medial occipital/middle cingulate cortices, respectively. Significant synchronization peaks in the time courses of C6 and C9 corresponded closely in time to peaks of the visual components (C1-C4), suggesting that C6 and C9 are responsive to similar stimuli.

## Discussion

The development of a data-driven approach for examination of task-driven brain responses requires a fundamental shift in both conceptual framework and analytic strategy. Here we demonstrate the utility and power of an inter-subject phase synchronization approach to describing task-driven brain responses. The spatial patterns and synchronization time courses of each network replicated across two independent samples of subjects. Importantly, the synchronization networks for each task spanned the majority of the brain’s cortical and sub-cortical regions, indicating that the task-response of the brain is not ‘localized’ to a single network or region, but in line with earlier predictions^45^, the majority of the brain responds in differing ways to task demands. The inter-subject phase synchronization approach goes beyond a traditional “top-down” GLM analysis and provides a principled way to determine ‘when’ and ‘where’ these task-driven responses occur during the course of an experiment. Below we discuss what the findings from the inter-subject synchronization approach applied to a motor, theory of mind and working memory task might imply for a dynamical theory of motor action, theory of mind and visual working memory processes, respectively.

### Dynamics of Motor/Action Processes

Applied to a simple motor mapping task, we found that the synchronization approach achieved precise localization of task-driven brain networks, with each network dedicated to the movement of a unique body part. In addition, the approach revealed ‘non-motor’ networks with temporal dynamics related to other events in the task. While the ‘visual cue’ reference function with the top-down GLM approach identified both visual and FPN as active during this period of the task, our approach revealed these two networks do not respond simultaneously to the task, but consecutively. The results revealed that the FPN significantly synchronized before the onset of each motor block. In addition, the spatial pattern of this network also included primary motor areas, indicating that these regions may serve as a preparatory process for motor/action control. This observation is further supported by previous findings of activation in the FPN and motor cortex during action preparation^46–48^. Synchronization of the FPN is followed by synchronization of the visual network, including areas of the posterior parietal cortex, that occur close in time to the visual cue, reflecting visuospatial processing of the visual cue indicating the appropriate motor movement. Together, these findings indicate a sequence of motor preparation, visuospatial processing, and motor action encompassing fronto-parietal, visual and motor areas of the brain that constitute a successful motor action. Importantly, this information regarding temporal dynamics of brain responses during task performance cannot be obtained using a traditional GLM approach.

### Dynamics of Theory of Mind Processes

The synchronization approach applied to the theory of mind task produced several visual and cognitive networks with complex temporal dynamics not predicted by the ‘top-down’ GLM approach. Only the bilateral fusiform/right TPJ network (C3) was related uniquely to a particular block of the task. Consistent with these brain areas role as a potential theory of mind ‘module’^38,39^, this network significantly synchronized only during the theory of mind block types. These areas were also identified by the subtraction contrast between the theory of mind and random blocks from the top-down GLM approach. However, examination of other network synchronization dynamics challenges the inference that these regions are selectively responsible for theory of mind processing. For example, the posterior parietal network (C4) exhibited sustained synchronization across the entirety of both task runs (theory of mind and random blocks), even during synchronization of the bilateral fusiform/tempo-parietal junction network (C3). This suggests that the posterior parietal network is a sustained attention process that biases the brain toward relevant sensory signals of the task, consistent with previous research^49,50^.

In addition to this network, a network (C1) consisting of the dACC, PCC and left temporal lobe, and a network (C6) consisting of the supplementary and left motor cortex, exhibited significant synchronization towards the latter end of all task blocks. The PCC and left temporal lobe are parts of the DMN^51,52^, which has been implicated in theory of mind processing^53,54^. The dACC is often considered a crucial component of the salience network^55^. Because these regions synchronize after the fusiform/TPJ network and synchronize in both random and theory of mind blocks, they may constitute a form of theory of mind processing important for the subsequent decision of intentional versus random interactive stimuli, as opposed to immediate processing of the stimuli in both blocks. Consistent with the requirement of a motor response to indicate the participant’s answer at the end of the task block, synchronization of the supplementary/left motor cortex may represent a preparatory motor signal for the subsequent motor response. Overall, these findings suggest that a single region or network (e.g., TPJ), as identified via a GLM analysis does not adequately capture the rich temporal dynamic involved in the theory of mind task. Rather, an inter-subject phase synchronization approach identifies a number of networks spanning many cortical structures with variable task-driven temporal dynamics across the course of the task.

### Dynamics of Visual Working Memory Processes

The synchronization approach applied to the visual N-back task with four categories of visual stimuli produced unique functional partitions of visual and posterior parietal areas. Four visual/parietal networks exhibited significant synchronization towards the end of each working-memory task block reflecting synchronized de-activations of these regions across subjects (**S5 Figure/Discussion**). This is consistent with evidence that both primary visual and higher-order visual areas are involved in the maintenance of visual representations for use in cognitive processing^56,57^. These networks encompassed several areas of the fusiform and visual/posterior parietal cortices that are often uniquely attributed to specialized processing of faces^58^, places^59^, human body^60^ or objects^61^. However, in accordance with evidence that encoding of visual features is distributed^62,63^, all four networks exhibited synchronization in response to all four stimuli type blocks. Rather than specialized encoding of faces, places, tools or body parts in a particular network, these networks seem to encode more basic features of visual stimuli that are present in all four stimuli types.

Examination of the other networks discovered through the synchronization approach reveals that primary visual and higher-order visual areas are not the only areas responsible for short-term maintenance of visual features. In fact, a number of networks, including the default mode network, the fronto-parietal network, basal ganglia, the dACC/PCC network observed in the theory of mind task, and a precuneus/DLPFC/TPJ network, exhibited synchronization to N-back task blocks. These findings suggest that a theory of visual working memory processes involving neural processing within a single brain network or brain region is inadequate. Rather, a comprehensive theory of working memory processing involving a coalition of fronto-parietal and visual networks provides a more comprehensive description of visual working memory processes.

### Dynamical Theory of Cognition and the Inter-Subject Synchronization Approach

Within a complex adaptive systems framework^64–66^ (Bolt, Anderson and Uddin, Under Review), the experimenter is interested in the temporal dynamics of the neural system as it evolves through time. The complex adaptive system framework compliments a dynamical theory of cognition where task-based psychological processes are described in terms of the temporal dynamics of brain networks as they synchronize simultaneously or sequentially over time. Conventional ‘top-down’ GLM approaches reduce this dynamical process into a dichotomous ‘active/non-active’ inference on a localized set of regions or networks. We argue that the inter-subject synchronization approach provides task fMRI researchers with the ability to recover traditional GLM activation patterns while also identifying novel low-frequency synchronization dynamic brain networks across time. Accordingly, the dynamical theory of cognition proposed in this study is a network-based theory of cognition, whereby psychological processes are constructed out of a dynamic coalition of brain networks^13,67^, as opposed to the fixed functions of a few brain regions or a single network. Consistent with arguments that psychological processes map onto a large range of functionally distinct brain networks^14,68^, the synchronization approach discovered a large number of networks involved in each task, covering a large expanse of cortical and sub-cortical brain structures.

### Limitations of the Inter-Subject Synchronization Approach

While synchronization provides a flexible exploratory approach for the study of task-driven brain responses, the approach does require certain assumptions to be met. First, the approach requires that the task run presented to each participant is identical in timing across all participants. This is essential to the approach, as the synchronization metric measures the extent of similar temporal dynamics in the time course across participants at each time point. Thus, task designs that utilize randomized presentation of task stimuli across participants would not be appropriate for the approach presented here, nor would tasks with counterbalanced task blocks across participants, unless the BOLD time courses are reordered before the analysis. Second, differences in hemodynamic responses and cerebral blood flow dynamics between participants would have adverse effects on the synchronization metric across participants, to the extent that these differences result in disparate timing in hemodynamic responses. In the traditional GLM reference function approach, these differences can be partially accounted for by additional regressors that model between-subject differences in hemodynamic durations and onset delays^69^. However, in the future, techniques accounting for these between-subject differences could be conceivably applied to the synchronization approach as well. As demonstrated by the current results, the approach still yields precise estimates despite these concerns.

## Conclusion

The goal of this study was to provide empirical support for a dynamical theory of cognitive processes through use of a methodological framework of synchronization dynamics that incorporates insights regarding task-driven signal variability into an exploratory task fMRI approach. The synchronization approach produced rich and varied insights into the dynamical neural interactions underlying the psychological processes involved in simple motor responses, as well as higher cognitive functions. Future studies are needed to develop the synchronization approach into a standardized methodological approach for analysis of task fMRI. For example, simulation studies could provide insights into the sample size and task designs necessary to achieve robust and replicable synchronization estimates. We believe that the dynamical study of cognitive processes made possible by the inter-subject synchronization approach implemented in the current study provides a valuable tool for cognitive neuroscientists.

## Acknowledgements

This work was supported by the National Institute of Mental Health [R01MH107549] and a NARSAD Young Investigator Award to LQU.

## Materials and Methods

### fMRI Data

Neuroimaging data from 150 unrelated, healthy, right-handed adults (Age Ranges: 22-25: 46, 26-30: 54, 31-35: 48, 36+: 2; 75 female) made available through the Human Connectome Project (HCP) 2014 release were used for this study. Seventy-five of these participants were used for the main results, and the other 75 participants were used as a confirmatory replication sample. Participants were recruited from the surrounding area of Washington University (St. Louis, MO). All participants gave informed consent before participating in the study, as described in Van Essen et al.^70^. Three task scans, a motor task, a theory of mind task, and working-memory task were used for analysis^35^.

### Task Descriptions

All details for the motor, theory of mind and working memory task used in this study are provided in^35^. The motor task was chosen because the task involved simple motor movements and the movement-brain activation mappings are well understood^36^, which provides an initial test case for the synchronization approach. In the task, participants are presented with visual cues that ask for movements of five specific body parts: tap left or right fingers, squeeze left or right toes, or move their tongue. The task involved two runs which both consisted of 13 movement blocks lasting 12s, preceded by a visual cue lasting 3s, and three fixation blocks lasting 15s. The reference functions used for the task were provided through the HCP, and included five regressors modeling the five specific movement types, and one regressor modeling the visual cue that indicated the start of each movement block. In addition, an all movement regressor was created that modeled all movement type blocks.

To test the synchronization approach on a task involving higher cognitive processing, the theory of mind and working-memory tasks were chosen. The theory of mind task involved higher-level processing of dynamic visual stimuli, consisting of short video clips of either interacting or randomly moving shapes, and rating periods where the participants indicated either of three choices: the objects had a social interaction, Not Sure, or No Interaction. The entire task included two runs, each including five 20s video blocks: 2 Interaction (theory of mind) and 3 Random for run one, and 3 Interaction and 2 Random for the run two; and five 15s fixation blocks. The reference functions used for this task were provided through the HCP, and included two regressors for interaction (theory of mind) video blocks and random video blocks, respectively (the rating period was not modeled). In addition, an all-block regressor was created that modeled both interaction and random video blocks together.

The working-memory task was a N-back task that involved visual processing of category specific visual stimuli. The entire task consisted of two runs, that each included ½ 0-back (a target visual cue is presented at the beginning of each block and the participant is directed to respond when that visual stimulus is presented) and ½ 2-back blocks (the participant is directed to respond when the current visual stimulus is identical to one presented two presentations back) that presented one of four stimulus types blocks (body parts, faces, places and tools). Thus, there were eight 25s task blocks for each run representing all possible combinations of task type (0-back and 2-Back) and stimulus type (body parts, faces, places and tools), along with four 15s visual fixation blocks. At the start of each block, there is a 2.5 sec visual cue indicating the task type (0-back versus 2-back), followed by ten 2.5s trials, where a stimulus was presented for 2s, followed by a 500ms inter-trial interval. The reference functions used for this task were provided by the HCP, and included eight regressors each modelling one of the eight possible task blocks (e.g. 0-back face block). In addition, four regressors were created modelling the four visual stimulus types (regardless of task type), and an all-block regressor modeling all task blocks.

### Data Preprocessing

Minimally preprocessed data provided through the HCP were used for further analyses. The minimal preprocessing pipeline involved gradient distortion correction, motion correction, registration to the Montreal Neurological Institute (MNI) template, and intensity normalization. The details of the minimal preprocessing pipeline are described in^71^. Additional preprocessing steps included demeaning, variance normalization (normalizing the data to their standard deviations from the mean, *z-score*) and concatenation of the time series from the right-left and left-right runs, respectively for each task using the Connectome Workbench^72^. Data was resampled to 3mm voxel size for computational feasibility, time courses were despiked using AFNI’s 3dDespike, an interpolative scrubbing procedure, and time-series were then detrended using FSL’s nonlinear Gaussian weighted least-squares fitting with a cutoff of 100s^17^. Of note, this high-pass filtering procedure only removes a part of the low-frequency signal below the cutoff of 100s and frequency bands can be still be studied below this cutoff, as demonstrated by other inter-subject correlation applications^73^. The functional data were then spatially smoothed (5mm full width at half maximum) with FSL, and nuisance covariate regression was performed (Friston’s 24 motion parameters, namely each of the 6 motion parameters of the current and preceding volume, plus each of these values squared, ventricle and white matter signals) using the Data Processing and Analysis for Brain Imaging (DPABI) toolbox^74^. No participants included in the analysis displayed gross motion (relative Root-Mean Squared-Framewise Displacement^75^; RMS-FD < 0.55mm;^76^.

### Standard General Linear Model Activation Estimates

Large-sample (n = 486) activation maps derived from the standard general linear model for each reference function were collected from NeuroVault^77^, to ensure that differences with the synchronization approach were not the result of insufficient statistical power. A conventional mixed-effects GLM analysis was conducted to derive activation estimates for each reference function, and is described in^35^.

### Determination of Frequency Band for Phase Analysis

Because the instantaneous phase synchronization approach is most effective for a band-pass limited signal^28^, we first conducted an eigenvalue synchrony analysis^29^ for all three tasks across frequency sub-bands to determine the frequency band that demonstrated the strongest average inter-subject synchrony. The eigenvalue synchrony analysis is a *static* inter-subject synchrony analysis, because it computes inter-subject synchrony across the course of the entire task scan. This approach allowed for a data-driven assessment of average synchrony in each frequency band to determine which band to use for the instantaneous phase synchronization analysis. Four frequency sub-bands were constructed through the stationary wavelet transform (SWT) as described by Kauppi et al.^73^ and implemented in the Inter-Subject Correlation (ISC) toolbox^32^. The SWT is more desirable over standard implementations of discrete wavelet transforms, as it is *time-invariant*, making the filter bank construction robust to small differences in hemodynamic lag between participants that could lead to large inconsistencies in the estimation of inter-subject synchronization. The SWT algorithm operates through successive splitting of low pass-filtered signals into new low- and high-pass signals^73^, and results in a logarithmic scale of the frequency sub-bands. We conducted a 4-scale SWT decomposition, resulting in 4 frequency sub-bands ranging from higher (Band 4) to lower frequency bands (Band 1). The approximate frequency characteristics, calculated as the frequency range in which 90% of spectral power is contained, of each band is: Band 1: 0.01 −0.085 Hz, Band 2: 0.08 −0.18 Hz, Band 3: 0.13 −0.4 Hz, Band 4: 0.35 −0.7 Hz. The periodogram of the four frequency bands in a sample time signal are displayed in **S2 Figure**. Of note, the SWT-filter is associated with a constant phase delay, and results in a constant 13 TR (6 sec) time delay for the filtered signal compared to the original signal. This time delay was compensated for by temporally shifting the standard GLM reference functions (discussed below).

The inter-subject eigenvalue synchrony approach was applied within each sub-band. The approach is conducted on a voxel-by-voxel basis and provides an assessment of group-wise synchrony in the time series across all subjects at each voxel. The strength of synchronization for each sub-band was computed by simply averaging the group-wise synchronization estimates across all voxels within the brain. The resulting eigenvalue maps (representing eigenvalues at each voxel in the brain) were thresholded using the permutation testing framework implemented by Kauppi et al.^32^, but adapted for constructing a null distribution of eigenvalues. For further details regarding the inter-subject eigenvalue synchrony analysis and permutation testing framework implemented in this study see **Supplemental Experimental Procedures**.

In addition to determination of the strongest synchronization sub-band, the group-wise static synchronization estimates at each voxel across frequency sub-bands were compared with the ‘activation’ estimates (z-score estimates) derived from the standard GLM approach using the conventional reference function. In particular, static synchronization estimates for the motor, theory of mind and working memory task were spatially correlated with the absolute-valued all motor movement, theory of mind, and 2-back activation maps, respectively. This allowed for an assessment of the degree of correspondence between the identification of task-driven brain responses between the static inter-subject synchrony analysis and the standard GLM approach.

### Instantaneous Phase Synchronization Calculation

The strongest synchronized frequency band, in terms of greatest average synchrony (i.e. eigenvalues) across the brain, was used for the instantaneous phase synchronization approach. Phase-based representations of fMRI signals have been used successfully in other fMRI analyses^25,78^, and has been previously validated for measurements of inter-subject synchronization by Glerean et al. (^28^. The instantaneous phase synchronization analysis works by first creating an analytic (i.e. complex-valued) representation of the original signal using a signal processing tool, known as the Hilbert transform, which proceeds in the following steps: 1) the Fourier transform of the signal *x(t)* is computed, 2) the negative frequencies are removed, and 3) the inverse Fourier transform is computed. The result is a complex-valued signal that can be represented as the product of *a(t)* the instantaneous envelope and φ(*t*) the instantaneous phase. Using the phase information from each participant’s signal, the normalized average angular distance^32^ between all participant-pairs can be computed as a group-wise measure of synchrony for each time point (TR) that varies from 0 to 1, where a value of 1 represents complete similarity of phase signals, and a value of 0 represents the complete absence of similarity of phase signals.

### Independent Component Analysis of Synchronization Time Series

The result of the instantaneous phase synchronization analysis is a time-series of synchronization values for each voxel in the brain, representing the average synchrony (the average absolute angular distance) across participants for each time point (TR). Rather than a conventional ROI-based analysis of synchronization, we chose to use a data-driven independent component analysis (ICA) that incorporates synchronization time signals across the entire brain to estimate possible synchronization networks that ‘pipe in’ and ‘fade out’ of the observed synchronization signals across the course of the task scan. The ICA was implemented through FSL’s MELODIC software^33^. This approach is equivalent to a single-subject ICA applied to group-level synchronization time series across all voxels in the brain, rather than the original signal time-courses of all voxels.

### Assessment of Number of Components and Replication

We estimated a range of lower model-order ICAs, including 10, 15 and 20 component solutions. Lower model-order ICAs were estimated for two reasons: 1) As most traditional task-fMRI analyses are concerned with a small subset of brain regions involved in any one task, a lower number of synchronization networks were estimated, and 2) the components are estimated from a ‘single subject’ or single brain of group-wise synchrony time series, which contains less data points (as opposed to group-level ICAs) for robust/replicable estimation of higher-order model ICAs. Of the 10, 15, and 20 component ICA solutions, we chose the solution that was most replicable in another confirmatory sample of 75 participants. ‘Replicability’ was measured in three ways: 1) the number of components that had ‘matching’ component pairs in the confirmatory sample in terms of visual examination between the original and confirmatory ICA solutions, 2) the spatial correlation between the ‘matching’ unthresholded component pairs from the original and confirmatory ICA solutions, and 3) the temporal correlation between the synchronization time series (computation described below) of the ‘matched’ components from the original and confirmatory ICA solutions. In addition to the determination of the number of components, the confirmatory sample also allowed an assessment of the degree to which the synchronization approach replicates across samples.

After ICA estimation of the chosen model-order, classification of components into noise and non-noise components was conducted in the same manner as ICA applied to normal BOLD time series: components with many strongly weighted voxels in white matter or cerebrospinal fluid (CSF) were classified as noise

### Component Synchronization Time Series and Statistical Significance

We implemented a bootstrap resampling procedure described in Glerean et al.^28^ to test for significant synchronization values (normalized averaged angular distance) at each time point over the course of the task scan for each component. For each task, the procedure involved bootstrap resampling of the original instantaneous phase time series (i.e. the phase representation of the original filtered time-series constructed through the Hilbert transform) to construct a null probability distribution estimated over 1000 permutations of each voxel’s time series of 568, 548 and 810 time points for the motor, theory of mind and working-memory task, respectively. Because a significance test is applied at every time point in the synchronization time course of each component, we implemented a multiple comparisons correction described in Alluri et al.^79^ that first estimated the number of independent time points in the time course, accounting for autocorrelation, and then applied a Bonferroni correction (α = 0.05 / # of independent time points).

The significance test could not be applied to the normalized component time courses derived from the ICA because they were not in the original scale of the phase synchrony metric (normalized average angular distance). Thus, the thresholded spatial components (threshold applied via mixture modeling;^80^ were used as weighted masks and applied to the original synchronization time courses. Specifically, the thresholded spatial components were used to extract the weighted mean time series from the synchronization time courses within each component mask, representing the weighted average synchrony time course for each component. The significance test was applied to the time points of the average synchrony time course of each component. Of note, the average synchrony time courses did not exhibit significant deviations from the normalized time courses derived from the ICA, as indicated by the temporal correlation between the two time courses (Motor: *r̄* = 0.96,*SD* = 0.04; TOM: *r̄* = 0.87,*SD* = 0.04; WM = *r̄* = 0.88,*SD* 0.06.

### GLM Reference Functions and Synchronization Comparison

As described above, we used the convolved block regressors of the GLM from the motor, theory of mind and working-memory tasks as reference functions for the synchronization time series. Reference functions represent the hypothesized BOLD activation for the cognitive/motor/affective process of interest. As described above, the SWT algorithm produced a positive 13 time-point (TR) shift in the low-frequency filtered BOLD signal (~ 0.01 to 0.085Hz) compared to the original signal. To compensate for this shift in the signal, all GLM reference functions were positively shifted by 13 TRs (6s) and 13 zero values inserted at the beginning of the time course to preserve the length of the signal. As noted by others^81^, conventional double-gamma HRF convolved regressors attempt to model a late undershoot of the HRF, which would not presumably be present in the case of a voxel *synchronized* to the task blocks, thus we used the gamma HRF to account for HRF lag and width, without an undershoot. The direct association between the synchronization time series and a chosen reference function was computed through the Pearson correlation between the two time courses.

### Comparison of Synchronization Components with Group ICA BOLD Components

An important question is the extent to which the functional connectivity relationships in the group-wise synchronization time series reflect functional connectivity relationships observed in the original BOLD data. ICA BOLD networks were estimated for all three task scans on the original sample of 75 participants using a temporal concatenation group ICA approach^82^. The temporal concatenation approach to group ICA involves the application of a single 2D ICA on the concatenated data matrix, created by vertically stacking 2D participant data matrices (*voxels/brain regions × time points*) along the time dimension. Because it wasn’t known at what resolution the synchronization networks are most likely to replicate in the group BOLD ICA solution, a lower-order (n = 10 components) and higher-order solution (n = 100) were computed. To detect a possible ‘match’ between the synchronization and BOLD components, spatial correlations were computed between the unthresholded synchronization components and each unthresholded BOLD component in both solutions (n = 10 and n = 100) for each task scan. A ‘match’ between a synchronization component and a BOLD component was defined as a strong spatial correlation (*r* > 0.4) between the *unthresholded* components and similar visual overlap between the *thresholded* components. These results are discussed in **S4 Figure/Discussion**.

### Comparison of Average BOLD Dynamics with Synchronization Estimates

Another question of interest is the relationship at any time point between average BOLD magnitude across participants, and the degree of synchronization observed across participants. Significant positive correlations between average BOLD time courses and synchronization time courses would indicate that periods of strong synchronization across participants occur during periods of increased BOLD activation across subjects. Because the synchronization metric was computed on filtered time-series, we applied the same SWT decomposition to construct four frequency sub-bands and used the BOLD signal sub-band (~0.01 – 0.085Hz) used in the synchronization approach. The filtered BOLD time course for each subject were then averaged across subjects to create an average BOLD time course at each voxel. The thresholded synchronization components were used to extract weighted average BOLD component time courses from the average BOLD signals. Because the comparison of interest was between BOLD *magnitude* changes and synchronization estimates, rather than increases or decreases in BOLD signal, the *absolute valued* time series was used to compared with the synchronization estimates. The association between the two time courses was computed through the Pearson correlation between the filtered average BOLD time course and the synchronization time course for each component. Because the number of components in each task were small and limited the number of possible BOLD-synchronization comparisons, we also computed correlations between the average (absolute valued) BOLD time courses and synchronization estimates from 264 ROIs (6mm spheres) as defined by Power et al.^83^. These results are discussed in **S5 Figure/Discussion**.

## Supplemental Materials and Methods

**S1 Figure.**
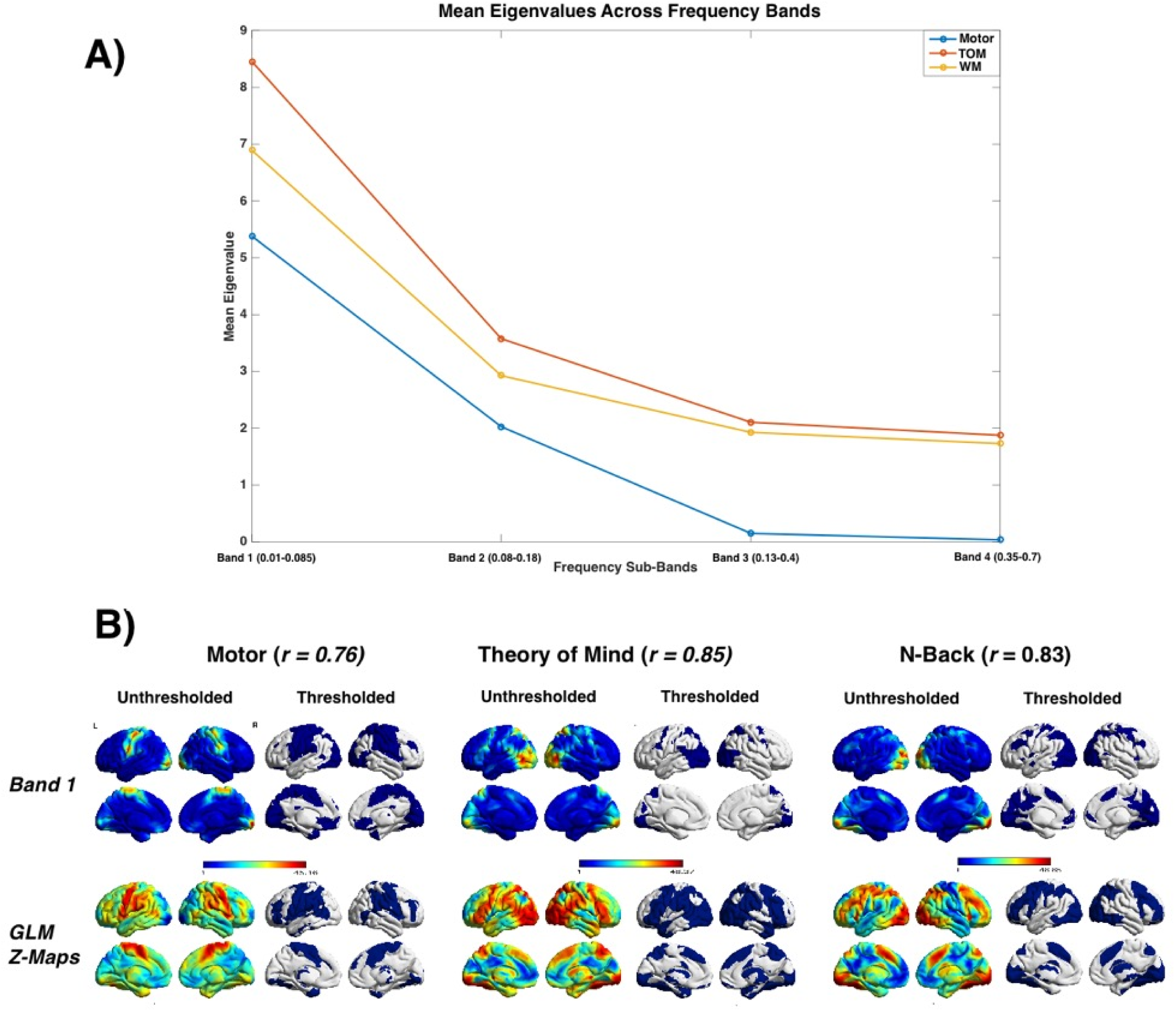
Static Inter-Subject Synchrony Across Frequency Sub-Bands and Comparison with GLM. (TOM = Theory of Mind; WM = Working-Memory; L = Left; R = Right). **A)** Mean eigenvalue (measure of static inter-subject synchrony) across the brain computed in Bands four, three, two and one for the motor (blue), theory of mind (red) and working-memory task (yellow). As can be seen from the plot, mean eigenvalues increase exponentially from Band four to Band one. **B)** Unthresholded and thresholded eigenvalue maps, visualized with BrainNet Viewer (27) for Band 1 and traditional GLM z-statistic activation/de-activation maps computed using a standard group-level analysis; unthresholded to the left, thresholded to the right: *z* > 3.1. For each task, the correlation between the Band 1 map (the largest correlation of all sub-bands for all tasks) and the absolute-valued z-statistic activation map is presented besides the task headings.

**S2 Figure.**
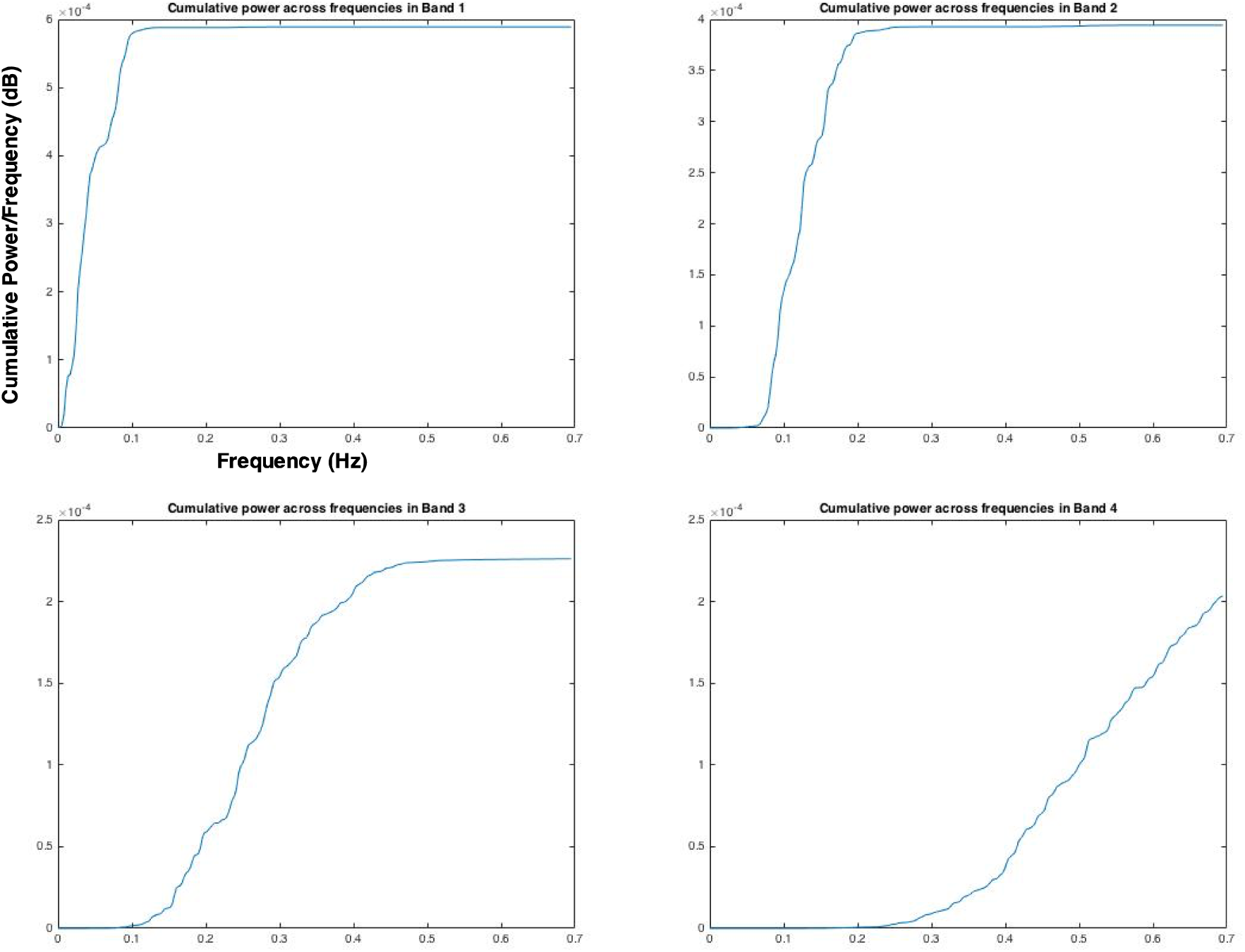
Cumulative Power Plots by Frequency Band. Cumulative distribution plots of the power spectral density by frequency for each of the four wavelet-constructed frequency bands.

**S3 Figure.**
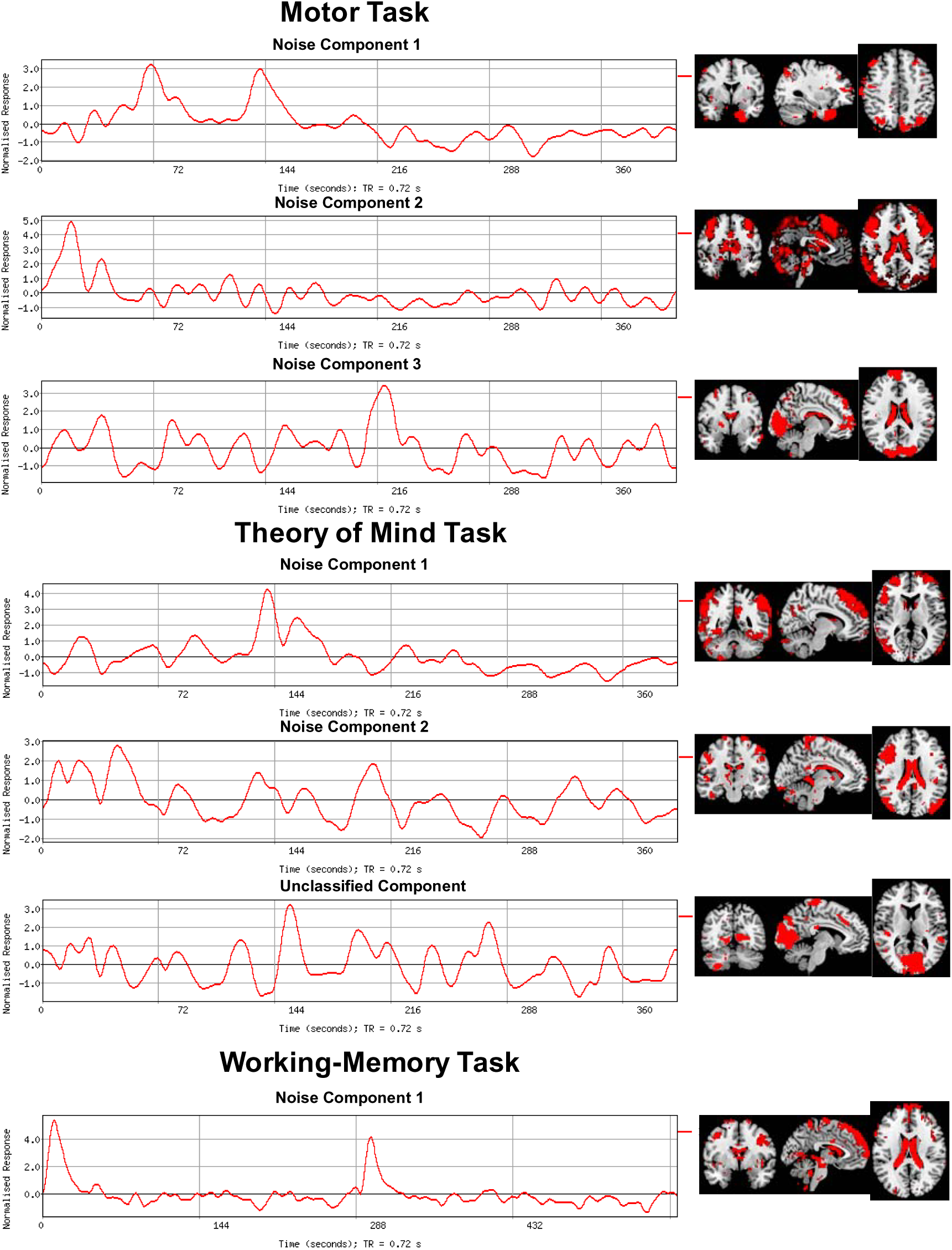
Noise Components and Unclassified Components in Motor, Theory of Mind and Working-Memory Tasks. Components in the ICA solutions that were classified as noise, based on voxel weight spatial patterns in white matter and cerebrospinal fluid regions. In the theory of mind task, a single component was not-interpreted (unclassified) because it was not discovered in a corresponding confirmatory sample.

**S4 Figure/Discussion.**
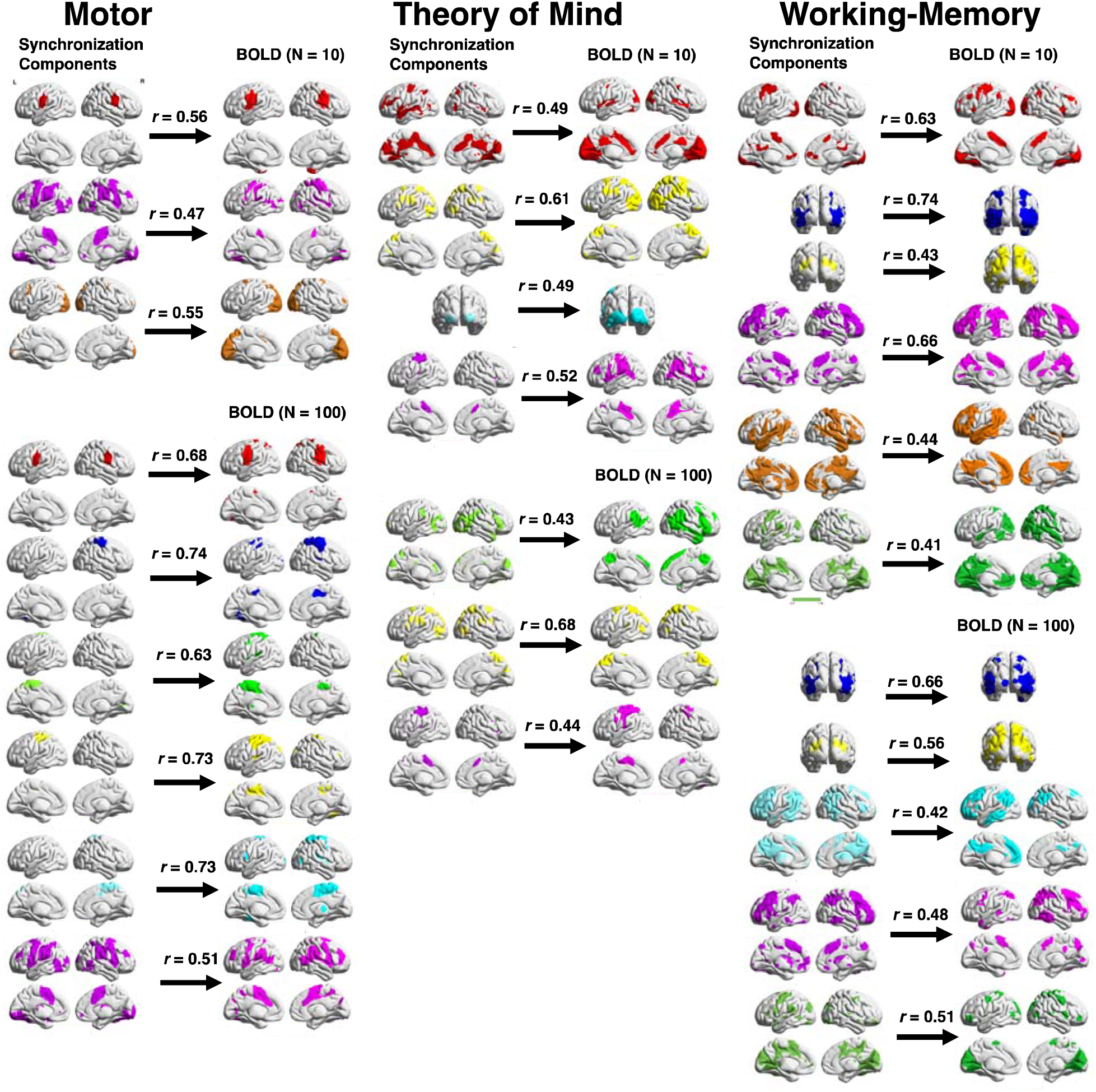
Pair-wise Comparison of Synchronization and Group-ICA Bold Components. Spatial weights and spatial correlation between synchronization components and matching component in the 10-component and 100-component group ICA solution derived from the BOLD time courses for each task. For each task column, synchronization components are visualized on the left and the corresponding group-ICA BOLD component on the right. Group-ICA component spatial weights were thresholded in the same manner as in the synchronization components^45^. The spatial correlation was computed between the unthresholded spatial weights of the matching components, and is represented above each arrow.

### Comparison of Functional Connectivity Relationships in Synchronization and BOLD Time Series

An important question is the extent that functional connectivity relationships in the group-wise synchronization time series, modeled as spatially independent components of synchronization dynamics, reflect functional connectivity relationships observed in the original BOLD data. Voxels with similar BOLD signals would be expected to exhibit similar group-wise synchronization signals, but it is also possible that the group-wise synchronization signals, calculated across subjects, reveals functional connectivity relationships not observed at the within-subject BOLD level. To assess this question, we applied a temporal concatenation group ICA, at a lower model-order (n = 10) and higher model-order (n =100) solution, to the low-frequency filtered BOLD signals (~0.01 - 0.085Hz) of the original sample of 75 participants for all three tasks. The ICA components from all tasks for both 10 and 100 component solutions were then compared with each synchronization component from each task (see *Experimental Procedures*). For all tasks, the majority of the synchronization components were also observed in either the 10- or 100-component group-ICA solution, suggesting that much of the functional connectivity relationships among voxels in their BOLD signals at the within-subject level are reflected in the group-wise synchronization signals (**Figure 6**). However, several components derived from the group-wise synchronization signals were not observed in the group-ICA BOLD components, indicating that novel task-driven functional connectivity patterns are revealed by the group-level synchronization approach.

**S5 Figure/Discussion.**
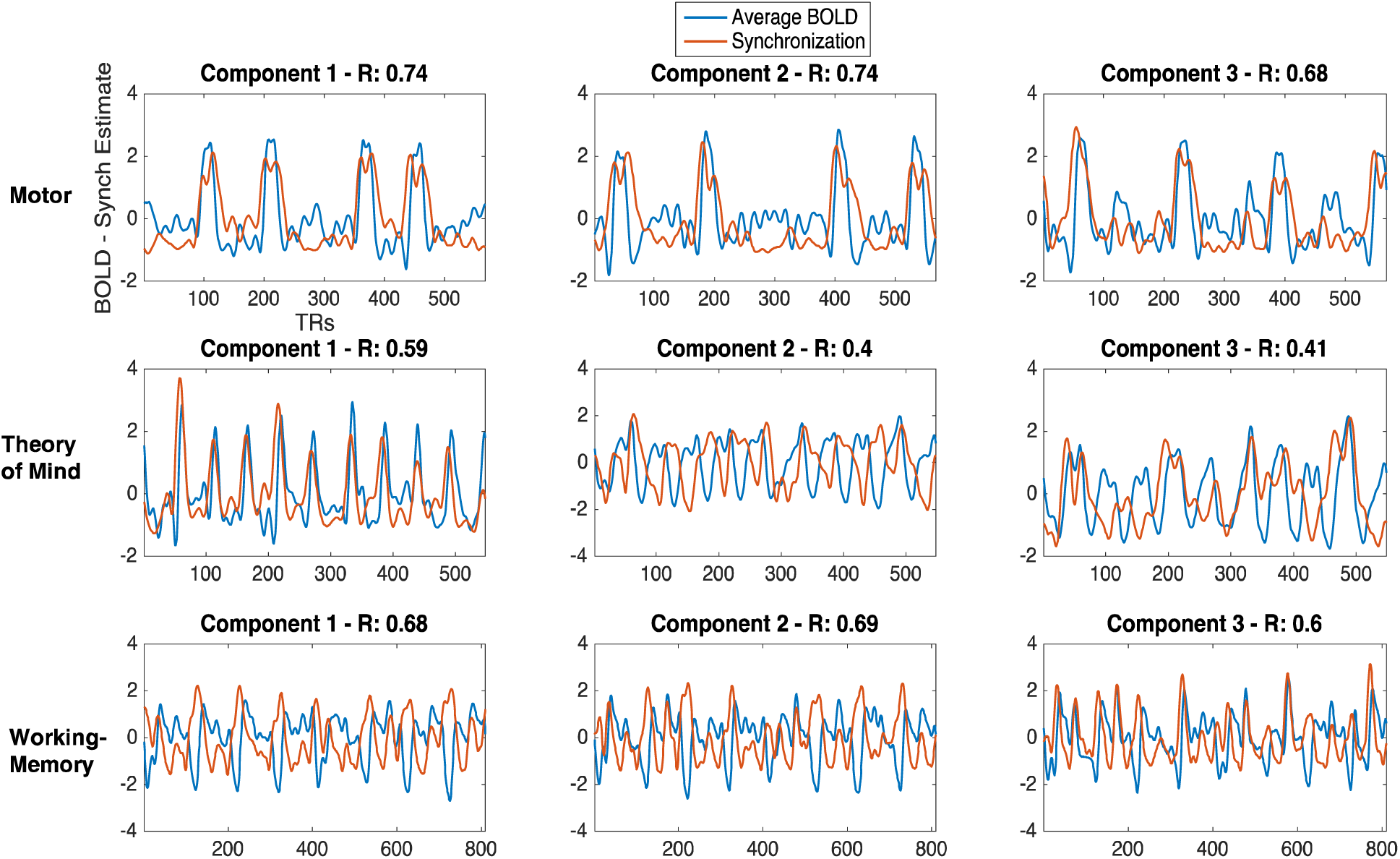
Average BOLD and Synchronization Comparison. Time courses of normalized (z-scored) average across-subject BOLD (blue) and group-wise synchronization estimates (orange) from the first three components of the motor, theory of mind and working-memory tasks. On top of each plot heading is the component number, and the correlation between the (absolute valued) average BOLD and group-wise synchronization time courses for that component.

### Average BOLD Dynamics and Synchronization Comparison

Another important question is the extent that group-wise synchronization estimates are associated with across-subject task-driven increases or decreases in BOLD signals. In particular, is low-frequency average BOLD magnitude across subjects associated with group-wise synchronization estimates across time points? For each component, the Pearson correlation between the low-frequency (~ 0.01 - 0.085Hz) average, absolute-value BOLD magnitude time course and synchronization time course was computed for all tasks. For all tasks, there was a moderate correlation between the component average BOLD magnitude and synchronization time courses (Motor: *r̄* = 0.68, *SD* = 0.08; TOM: *r̄* = 0.43, *SD* = 0.16; Working-Memory: *r̄* = 0.59, *SD* = 0.09). Because the number of components limited the number of possible BOLD magnitude-synchronization comparisons, Pearson correlations between average BOLD magnitude and synchronization time courses were also computed for 264 ROIs across the brain^46^. In agreement with the component results, there was a moderate correlation between ROI BOLD magnitude and synchronization time courses (Motor: *r̄* = 0.44, *SD* = 0.17; Theory of Mind: *r̄* = 0.41, *SD* = 0.14; Working-Memory: *r̄* = 0.45, *SD* = 0.13).

Visual comparison of the average across-subject BOLD and the synchronization time courses from the first three components of each task (**Figure 7**) illustrates the relationship between these two signals. The BOLD and synchronization signals from the motor components exhibit similar temporal dynamics during large positive amplitude BOLD spikes in the data, corresponding to the onset of motor trials. The synchronization time courses of C1 from the theory of mind task, and C1 and C2 from the working-memory task also exhibit strong peaks in synchronization during large positive and negative amplitude bold spikes, respectively. However, in time series with minimal BOLD amplitude spikes, such as the BOLD time courses of C2 and C3 from the theory of mind task, or low-BOLD amplitude periods in-between BOLD amplitude spikes, the correspondence between BOLD amplitude and synchronization time courses is reduced.

Studies of phase synchronization between broadband signals^67,68^, as the one used in this study (~0.01 to 0.085 Hz; this frequency range exhibited the strongest average synchrony across subjects), have demonstrated that synchronization estimates are more strongly correlated with synchronous large-amplitude signals spikes compared with lower-amplitude signal periods. This may explain the moderate relationship between average BOLD amplitude and group-wise synchronization estimates (*r* = ~ 0.3 - 0.7). This property of broadband phase synchronization is beneficial as an exploratory approach, as synchronous high-amplitude BOLD spikes in the data are expected to be of more task-relevance compared with low-amplitude signal periods. In addition, a restricted broadband signal in the frequency range of 0.01 to 0.085 Hz was chosen to ensure that all possible task-driven signal dynamics within this commonly studied frequency range are detectable by the phase synchronization approach. As demonstrated in the results for all three tasks, this signal adequately captures task-driven signal dynamics.

## Supplemental Experimental Procedures

The stationary wavelet transform (SWT) was implemented in the Inter-Subject Correlation (ISC) toolbox (21). The SWT is more desirable over standard implementations of discrete wavelet transforms, as it is *time-invariant*, making the filter bank construction robust to small differences in hemodynamic lag between participants that could lead to large inconsistencies in the estimation of inter-subject synchronization. The SWT algorithm operates through successive splitting of low pass-filtered signals into new low- and high-pass signals (40), and results in a logarithmic scale of the frequency sub-bands. We conducted a 4-scale SWT decomposition, resulting in 4 frequency sub-bands ranging from higher (Band 4) to lower frequency bands (Band 1). The approximate frequency characteristics, calculated as the frequency range in which 90% of spectral power is contained, of each band is: Band 1: 0.01 - 0.085 Hz, Band 2: 0.08 −0.18 Hz, Band 3: 0.13 −0.4 Hz, Band 4: 0.35 −0.7 Hz. The periodogram of the four frequency bands in a sample time signal are displayed in **Figure S2**. Of note, the SWT-filter is associated with a constant phase delay, and results in a constant 13 TR (6 sec) time delay for the filtered signal compared to the original signal. This time delay was compensated for by temporally shifting the standard GLM reference functions (discussed below).

The inter-subject eigenvalue synchrony approach was applied within each sub-band. The approach is conducted on a voxel-by-voxel basis and provides an assessment of group-wise synchrony in the time series across all subjects at each voxel. The eigenvalue synchrony analysis procedure is as follows: first, the pair-wise similarity between each pair of participants is calculated for every *voxel* in the brain (whole-brain *voxel-by-voxel* approach) by calculating the *Pearson product-moment correlation* (i.e. Pearson’s r) between each participant’s band-limited time series (calculated across all four sub-bands) at that voxel. The results of the analysis produce pair-wise correlation coefficients between all pairs of 75 participants at each voxel (this can be visualized in terms of a 75*75 participant-by-participant correlation matrix). The traditional approach (19, 20, 40) to calculating a group-wise synchrony estimate for each voxel, is to simply average across all pair-wise *z-*transformed correlation coefficients (essentially average the upper or lower triangle of the 75*75 participant correlation matrix). However, a more statistically principled way of estimating group-wise synchrony at each voxel that works over all participants simultaneously, and scales over sample size, would be to conduct an eigenvalue approach that estimates the first principle eigenvector and its associated eigenvalue through a singular value decomposition (SVD) of the participant-by-participant correlation matrix (18). The resulting eigenvalue is a measure of group-wise subject synchrony for that voxel, and ranges from 1 to the number of participants. The SVD algorithm provided through MATLAB was used to conduct this analysis. The strength of synchronization for each sub-band was computed by simply averaging the group-wise synchronization estimates across all voxels within the brain.

The resulting eigenvalue maps (representing eigenvalues at each voxel in the brain) were thresholded using the permutation testing framework implemented by Kauppi et al. (21), but adapted for constructing a null distribution of eigenvalues. Briefly, each participant’s band-limited time series is randomly circularly shifted so that participant time series are no longer aligned in time, and the first principle eigenvalue is computed, and this procedure is repeated 100,000 times to construct a null distribution of eigenvalues for each frequency band. P-values are computed for each voxel and then corrected for multiple comparisons using the false discovery rate (FDR; Benjamini and Hochberg, 1995).

